# Detection and molecular characterisation of SARS-CoV-2 in farmed mink (*Neovision vision*) in Poland

**DOI:** 10.1101/2020.12.24.422670

**Authors:** Lukasz Rabalski, Maciej Kosinski, Teemu Smura, Kirsi Aaltonen, Ravi Kant, Tarja Sironen, Boguslaw Szewczyk, Maciej Grzybek

**Affiliations:** Laboratory of Recombinant Vaccines, Intercollegiate Faculty of Biotechnology of University of Gdansk and Medical University of Gdansk, Abrahama 58, 80-307, Gdansk, Poland; Department of Virology, University of Helsinki, Haartmaninkatu 3, Helsinki FI-00290, Finland; Department of Veterinary Biosciences, University of Helsinki, Agnes Sjöbergin katu 2, Helsinki FI-00790, Finland; Department of Tropical Parasitology, Institute of Maritime and Tropical Medicine, Medical University of Gdansk, Powstania Styczniowego 9B, 81-519, Gdynia, Poland

**Keywords:** SARS-CoV-2, interstitial pneumonia, mink, transmission, spillover, zoonoses

## Abstract

SARS-CoV-2 is the aetiological agent of COVID-19 disease and has been spreading worldwide since December 2019. The virus has been shown to infect different animal species under experimental conditions. Also, minks have been found to be susceptible to SARS-CoV-2 infection in fur farms in Europe and the USA. Here we investigated 91 individual minks from a farm located in Northern Poland. Using RT-PCR, antigen detection and NGS, we confirmed 15 animals positive for SARS-CoV-2. The result was verified by sequencing of full viral genomes, confirming SARS-CoV-2 infection in Polish mink. Country-scale monitoring conducted by veterinary inspection so far has not detected the presence of SARS-CoV-2 on other mink farms. Taking into consideration that Poland has a high level of positive diagnostic tests among its population, there is a high risk that more Polish mink farms become a source for SARS-CoV-2. Findings reported here and from other fur producing countries urge the assessment of SARS-CoV-2 prevalence in animals bred in Polish fur farms.

## 1. INTRODUCTION

The identification of possible pathogen hosts and the study of transmission dynamics in their populations are crucial steps in controlling zoonotic diseases. In the case of SARS-CoV-2, the origin is most likely bats (Zhou et al., 2020), but the intermediate host has not been confirmed yet. SARS-CoV-2 seems to readily jump from people to other animal species, in particular carnivores, (Jo et al., 2020) raising the concern of new animal sources of COVID-19.

SARS-CoV-2 infections in mink have been reported from farms in Denmark, Netherlands, USA, Italy, Spain, Sweden, Greece and Lithuania (Hammer et al., 2021; Koopmans, 2020; Oreshkova et al., 2020; Oude Munnink et al., 2020) (Figure 1).

**Figure 1.**
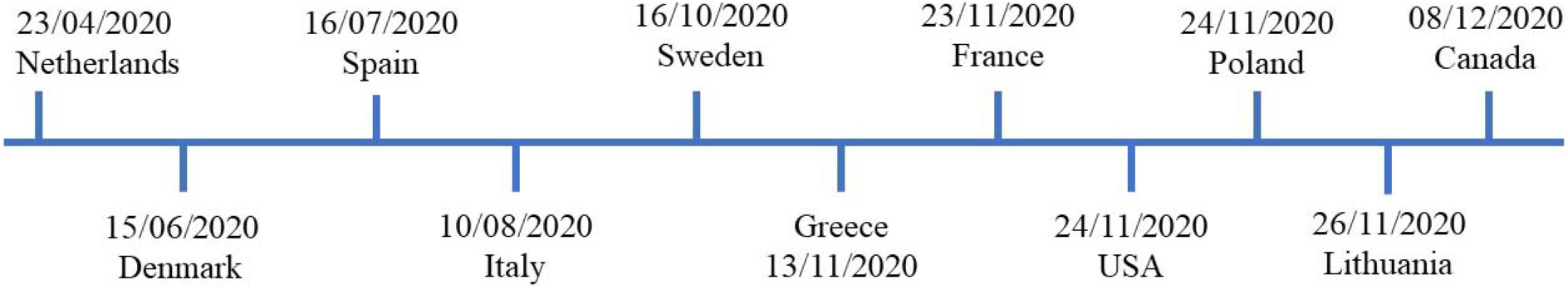
The time-line of SARS-CoV-2 infections in mink farms around the world, according to the World Organisation for Animal Health (World Organisation for Animal Health, 2020).

Due to SARS-CoV-2 outbreaks in mink farms and its appearance in the surrounding communities, the European Centre for Disease Prevention and Control and the WHO have emphasised the importance of surveying the host-animal interface and collaboration among virologists and epidemiologists to track and characterise viral mutations (WHO, 2020). Following SARS-CoV-2 infections in minks in the Netherlands, the Dutch Ministry of Agriculture decided to cull all minks from different farms. The Danish National Institute of Public Health announced the culling of all 17 million mink in the country after it found that the virus had spilt back from mink farms into the human community. Recent data available from Denmark and the Netherlands on these mink-associated SARS-CoV-2 variants suggest that these variants can spread rapidly in mink farms and among nearby human communities. Humans infected with the mink-related variants do not appear to have more severe clinical symptoms than those infected with non-mink-related variants (Oude Munnink et al., 2020).

After Denmark, Poland is the second-largest producer of mink pelts in Europe. The total number of fur animal farms in Poland is 810 (including fox, mink, raccoon dog, and chinchillas). There are 354 active Polish mink farms with ~6.3 million individual mink. In 2019, Polish mink farmers sold 8.5 million mink pelts (EU Fur Association, 2020; ZPP, 2020).

As of this writing, Poland has recorded over 1 135 676 COVID-19 cases, with over 22 864 total deaths (19/12/2020 ECDC, 2020).

Considering the recent reports of SARS-CoV-2 mink infections in other European countries and the high incidence of SARS-CoV-2 human infections in Poland, we conducted SARS-CoV-2 monitoring in mink on one farm located in Pomorskie Voivodeship in North Poland.

## 2. MATERIALS AND METHODS

### 2.1. Material collection

Throat swabs (BIOCOMA, Shenzhen, China) were collected from 91 minks from a mink farm located in Pomorskie Voivodeship in Northern Poland on 17^th^ November 2020. The material was collected from minks culled for pelting. The farm owner reported no respiratory symptoms in the animals.

### 2.2. Statistical analysis

The 95% confidence limits were calculated using bespoke software “PERCENTAGE CONFIDENCE LIMITS VS 13” (courtesy of Dr F.S. Gilbert and Prof. J.M. Behnke, University of Nottingham), based on the statistical tables (Sokal and Rohlf, 1995)

### 2.3. RNA isolation

A total of 150 μl of each sample from the swab in inactivation buffer was added to 300 μl of RLT lysis buffer (RNeasy Mini kit, Qiagen, Hilden, Germany). Samples were mixed by vortexing and incubated for 10 minutes at room temperature. After incubation, 400 μl of 70% ethanol was added to each sample and mixed by pipetting. The lysate was transferred to an RNeasy Mini spin column with collection tube and centrifuged 1 minute at 13 000 RPM. Columns were washed once with 700 μl RW1 and twice with 500 μl RPE. Between every wash, columns were centrifuged, and flow-through was discarded. Elution was performed by adding 50 μl of PCR-grade water to the column and incubating for 2 minutes. Columns were placed into new tubes and centrifuged at 13 000 RPM for 1 minute. After isolation, samples were stored for less than 2 hours at 4 ^◦^C before further steps. No human origin samples were processed at the same time.

### 2.4. Real-time RT-PCR

For each sample, the reaction mixture was prepared using a polymerase, water and primers and probes (Corman et al., 2020) in white 8-well q-PCR strips with optical clear caps according to Table 1.

**Table 1.** Composition of RT-qPCR reaction.

|  |  |
| --- | --- |
| qPCR grade water | 8 µl |
| 4x TaqPath 1step RT-PCR polymerase | 5 µl |
| Forward primer RdRp gene | 1 µl |
| Reverse primer RdRp gene |  |
| RdRp gene probe (FAM) |  |
| Forward primer E gene | 1 µl |
| Reverse primer E gene |  |
| E gene probe (HEX) |  |
| Sample | 5 µl |
| Volume of reaction | 20 µl |

Positive control and no template control (NTC) reactions were also prepared. Reactions were mixed and loaded into Light Cycler 480 (Applied Biosystems, Foster City, California, United States) with the program described below (Table 2). After each cycle of amplification, the signal from each sample was measured in both FAM (Rdrp gene) and HEX (E gene) channels.

**Table 2.** The program used for RT-qPCR.

| Step | No. of cycles | Temperature | Time |
| --- | --- | --- | --- |
| UNG incubation | 1 | 25 °C | 2 minutes |
| RT incubation | 1 | 50 °C | 15 minutes |
| Enzyme activation | 1 | 95 °C | 2 minutes |
| Amplification | 40 | 95 °C | 3 seconds |
|  |  | 60 °C | 30 seconds |

### 2.5. SARS-CoV-2 antigen detection in mink

We performed two different antigen tests to confirm the presence of virus antigen either in the swab or the serum samples.

#### 2.5.1. Antigen test 1 for the swab samples

Antigen tests were conducted on samples positive in qPCR. Three negative samples were used as control. A total of 150 μl of transport medium from each swab was transferred to tubes from a COVID-19 Antigen Detection Kit (Zhuhai Lituo Biotechnology CO., LTD.) containing an extraction buffer. Samples were mixed and incubated for 1 minute. Two drops of each sample were added to the sample window on test cassettes. Results were read after 12 minutes.

#### 2.5.2. Antigen test 2 for the serum samples

Ninety-one mink serum samples were tested with SARS-CoV-2 antigen ELISA (COV-04-S, Salofa Oy, Finland) according to the kit instructions. This test is a double-antibody sandwich ELISA. The results were obtained according to the formula based on the concentration standards provided in the kit. The cut-off value for this test was given as 2.97 pg/ml. The tests were repeated twice, and additional dilutions were performed to determine the final concentration as suggested in the kit instructions.

### 2.6. Full SARS-CoV-2 genome sequencing and classification

SARS-CoV-2 genome sequencing was performed at the Medical University of Gdansk, University of Gdansk, Poland and the University of Helsinki, Finland using samples containing RNA isolated from positive swabs (amplification of two target genes in RT-PCR) or inconclusive (only single target gene amplification). At Gdansk two independent protocols were used for SARS-CoV-2 genome sequencing: Illumina RNA prep with enrichment for respiratory virus oligos panel V2 followed by Illumina MiniSeq medium output run that produced 150-nucleotide paired-end reads, and ARTICv3 amplicon generation followed by Oxford Nanopore Technology MinION run (Quick 2020). No human origin samples were processed at the same time. No DNA / rRNA depletion methods were used. Reads were basecalled, debarcoded and trimmed to delete adapter, barcode and PCR primer sequences. The fasta files generated by the Illumina procedure were further analysed in Kraken2 software to classify every read to reference database containing viral and American mink genomes (Wood et al., 2019).

In Helsinki, the sequencing libraries were prepared using Illumina DNA prep kit (New England BioLabs). The library fragment sizes were measured using agarose gel electrophoresis and the concentrations using Qubit dsDNA HS Assay Kit (Life Technologies, Carlsbad, California, USA) and NEBNext Library Quant Kit for Illumina (New England BioLabs, Ipswich, Massachusetts, USA). Sequencing was conducted using MiSeq V3 reagent kit with 250 bp reads. Raw sequence reads were trimmed and low quality (quality score <30) and short (<50 nt) sequences removed using Trimmomatic (Bolger et al., 2014). The trimmed sequence reads were assembled against SARS-CoV-2 reference sequence (NC_045512.2) using BWA-MEM algorithm (Li 2013) implemented in SAMTools version 1.8 (Li et al., 2009).

### 2.7. Phylogenetic analysis of SARS-CoV-2 isolates

The dataset consisted of all genetic sequences of SARS-CoV-2 from this research, Poland, Germany, Lithuania, Latvia, Estonia, Russia, and Ukraine completed with representative pool “Europe” by Nexstrain.org (https://nextstrain.org/ncov/europe) resulting in a total number of 5778 entries. Phylogenetic analysis was performed using the procedure recommended by Nextstrain.org with modifications in subsampling region filtering procedure, where the number of sequences per country was 40 (Hadfield et al., 2018). In short: the Augur toolkit v10.1.1 was used for phylogenetic analysis and Auspice v2.10.1 for visualisation. Possible time of divergence for samples was inferred using the TreeTime pipeline implemented in Nextsrain analysis and presented on the phylogenetic tree (Sagulenko et al., 2018).

### 2.8. Ethical Approval

This study was carried out with due regard for the principles required by the European Union and the Polish Law on Animal Protection. No permit from Local Bioethical Committee for Animal Experimentation was obtained because animals were culled by the owner for production of pelts. Samples were collected *post mortem.*

## 3. RESULTS

### 3.1. Prevalence of SARS-CoV-2

Using two targets RT-PCR, antigen detection and NGS, we confirmed 15 SARS-CoV-2 positive individuals (16.5% [8.4-28.6]). Out of the 91 samples, 17 showed a positive RT-PCR result in the SARS-CoV-2 E gene amplification (18.7% [10.3-31.1]). Eight of these were also positive for the RdRp gene (8.8% [3.5-19.3]).

Table 3 summarises the results for the applied diagnostic approach. Highlighted samples are considered SARS-CoV-2 positive.

**Table 3.** Results of different techniques that confirm the detection of SARS-CoV-2 in samples collected from farmed minks in Poland. Column 1 - individual sample name; column 2 and 3 - RT-qPCR assay; column 4 - antigen detection assay in swab; column 5 - antigen detection assay in serum; column 6 - full viral genome sequencing. Highlighted samples are considered positive for SARS-Cov-2 by nucleic acid amplification technique, serology or NGS. Cp-crossing point, RdRp - RNA dependent RNA polymerase, E - Envelope small membrane protein, N - Nucleoprotein, sp - strong positive, p - positive, n - negative, nc - not checked, NTC - no template control.

| Sample Name | Cp for gene RdRp | Cp for gene E | The antigen in swab<br>(strong pos/pos/neg) | The antigen in serum<br>(concentration of N protein) | Sequence obtained<br>(full/partial/nc) |
| --- | --- | --- | --- | --- | --- |
| mink_4 | - | 24.90 | sp | 48.68 | full |
| mink_5 | 24.36 | 16.88 | sp | 3.3 | full |
| mink_6 | - | 34.88 | n |  | partial |
| mink_20 | - | - | p | 132.24 | partial |
| mink_27 | - | 35.05 | n |  | partial |
| mink_36 | - | - | p | 271.3 | partial |
| mink_39 | - | - | n | 35.59 | partial |
| mink_42 | - | 25.62 | p |  | full |
| mink_46 | 34.93 | 22.75 | p |  | full |
| mink_48 | 29.89 | 22.52 | sp |  | full |
| mink_49 | - | 27.79 | p |  | full |
| mink_50 | 32.39 | 21.61 | sp |  | full |
| mink_63 | - | - | n | 19.3 | partial |
| mink_67 | 36.88 | 23.34 | p |  | full |
| mink_70 | - | 24.40 | p |  | partial |
| mink_76 | 32.53 | 22.73 | p |  | full |
| mink_77 | 32.87 | 20.3 | sp |  | full |
| mink_83 | 15.89 | 21.27 | sp | 91.57 | full |
| mink_88 | - | 33.99 | p |  | full |
| mink_89 | - | 35.39 | n |  | nc |
| mink_90 | - | 35.61 | n |  | nc |
| NTC | - | - |  |  |  |
| Positive control | 16.91 | 23.31 |  |  |  |

### 3.2. SARS-CoV-2 antigens detected in mink

#### 3.2.1. Antigen test from the swab

Samples mink_4, mink_5, mink_48, mink_50, mink_77 and mink_83 gave highly visible signals in both control and test lines. Samples mink_20, mink_36, mink_42, mink_46, mink_49, mink_67, mink_76 and mink_88 gave next to a highly visible control line a much less pronounced test line. In all other samples, only the test line was visible. All eight real-time RT-PCR positive samples were also positive in the antigen test. Additionally, five E gene-positive samples were also positive in the antigen test, but four were negative. In the sample mink_20 that was positive in the antigen test, SARS-CoV-2 RNA was not detected by RT-PCR.

#### 3.2.2. Reads classification and SARS-CoV-2 genome sequences

The final validation of the detection of SARS-CoV-2 in minks was a classification of next-generation sequencing reads to the database containing reference viral, human and American mink genomes. Three independent approaches were used to achieve full virus genomes, and the data resulting from procedures are presented in Table 4. Only samples that gave rise to a complete SARS-CoV-2 genome sequence are shown. The number of Illumina reads generated for samples mink_4, mink_42, mink_49, mink_76 and mink_88 were not enough to produce full SARS-CoV-2 genomes. For these samples, genomes were obtained by ARTIC procedure.

**Table 4.** Total mean coverage (ONT and Illumina reads) for every sample that full SARS-CoV-2 genome was established. Last three columns represent how many Illumina reads were generated for each sample and how many of them were classified to SARS-CoV-2 or American mink genomes.

| Sample name | Mean coverage | Total Illumina reads | SARS-CoV-2 reads | Mink reads |
| --- | --- | --- | --- | --- |
| mink_4 | 949.6 | 152609 | 37 | 131204 |
| mink_5 | 2776 | 396259 | 159484 | 184135 |
| mink_42 | 847.8 | 66515 | 868 | 34968 |
| mink_46 | 1039.5 | 387682 | 25975 | 241788 |
| mink_48 | 1151 | 125328 | 9526 | 64363 |
| mink_49 | 394.6 | 487771 | 1711 | 348017 |
| mink_50 | 1839.6 | 425333 | 79589 | 246546 |
| mink_67 | 882.1 | 167963 | 10834 | 114163 |
| mink_76 | 723 | 99660 | 2105 | 77945 |
| mink_77 | 16078.5 | 2424311 | 1690172 | 662381 |
| mink_83 | 111978.9 | 13495934 | 12730431 | 611926 |
| mink_88 | 1205.2 | 1071003 | 2822 | 611787 |

#### 3.2.3. Phylogenetic analysis of SARS-CoV-2 in Polish farmed mink

The 12 mink originated SARS-CoV-2 sequences were checked for the presence of mink specific mutations detected earlier in minks from the Netherlands and Denmark, but none were found, pointing towards a recent introduction of SARS-CoV2 in Polish minks (Figure 2).

**Figure 2.**
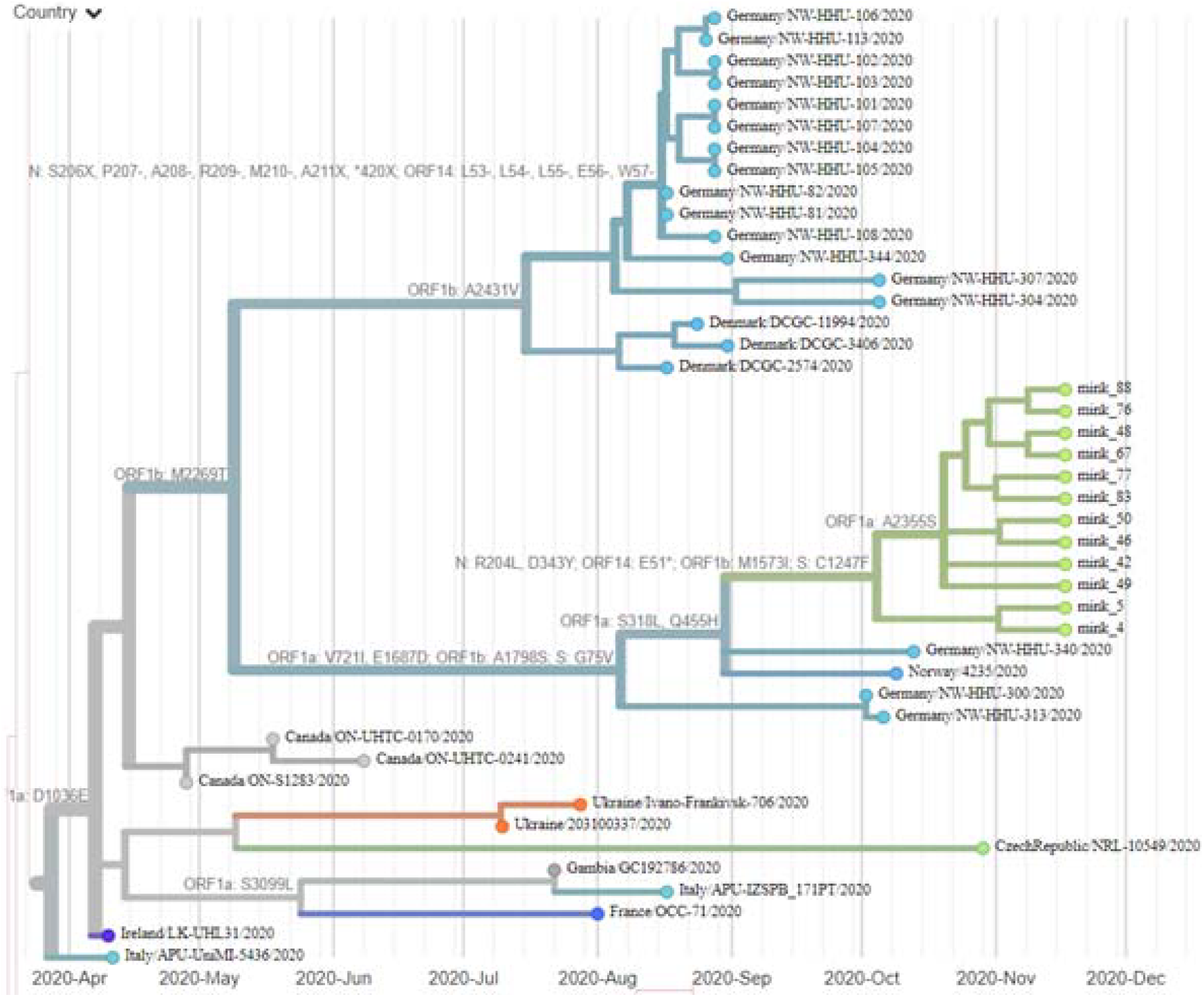
Phylogenetic tree estimating divergence time presenting only closely related isolates that were included in the dataset. Indications on branches represent amino acid changes that were associated with divergence. Produced using Nextstrain.org.

The alignment of the full genome sequence from 12 individual samples showed multiple polymorphisms on different nucleotide sites. Many of them gave rise to changes in amino acid in comparison to the reference (MN908947). Two specific mutations present in all samples were found in the S protein: G75V and C1247F. These mutations are present in 199 for S: G75V and 83 for S: C1247F isolates previously published in GISAID. Other rear amino acid variants present in every Polish mink SARS-CoV-2 isolate were found in five more proteins: nsp2, nsp3, nsp14, nsp15 and N.

Based on the dataset (Material and methods 2.6.) we inferred phylogenetic relationship by estimating divergence times between every isolate (Figure 1). The analysis estimates that the most recent common ancestor for Polish minks SARS-CoV-2 and two most similar sequences (German/NW-HHU-340/2020 and Norway/4235/2020) diverged around 31^st^ September 2020. Multiple mutations in amino acid sequences were recognised. If the molecular evolution starts after the virus introduction to the farm, this incident is estimated to happen around 4th October 2020.

## 4. DISCUSSION

Identifying new animal species that can serve as animal sources of SARS-CoV-2 and predicting where novel outbreaks are most likely to occur are crucial steps for preventing and minimising the extent of SARS-CoV-2 infections among humans (Hemida and Ba Abduallah, 2020). Recent reports confirmed the presence of SARS-CoV-2 in different animal species, including fur animals (i.e. minks and racoon dogs) (Abdel-Moneim and Abdelwhab, 2020; Freuling et al., 2020).

Here we report the presence of SARS-CoV-2 in minks from a fur farm in Northern Poland. We report a 16.5% prevalence of SARS-CoV-2 in the tested minks and confirm the presence of SARS-CoV-2 in farmed minks in Poland.

Poland is one of the largest fur producers in Europe. Considering the number of farmed minks in the country and the significant number of people employed in this sector, we seek to raise awareness in the scientific community and the mink industry that minks are susceptible to SARS-CoV-2 infection.

Previous studies reported the detection of viral RNA in airborne inhalable dust in mink farms (Oreshkova et al., 2020). Moreover, close contact of farmworkers with animals during feeding, culling, and dehiding increase the risk of exposure.

We believe that a country-scale biomonitoring programme should be activated as soon as possible to prevent the fur production sector from being a reservoir for future spillover of SARS-CoV-2 to humans. Samples for molecular diagnostics should be obtained for all farms in Poland following the highest standards of material collection, sample handling, and molecular detection of SARS-CoV-2.

We report a possible rise of the new genotype that possesses sporadic mutations through the full genome sequence. Two mutations located in Spike protein (G75V and C1247F) are present in all Polish mink virus isolates. G75V mutation is localised in the NTD domain and could be responsible for interaction with host receptors or stabilising Spike protein in a constrained prefusion state (Arya et al., 2021). Non-other sequences of SARS-CoV-2 deposited to date in GISAID possess these two mutations simultaneously (Shu and McCauley, 2017). For now, we do not know if this possible genotype has the ability for transmission to humans. Wide monitoring of humans living in close surroundings to the mink farm should answer this question.

## Data availability

The complete genome sequences of SARS-CoV-2 isolated from farmed Polish minks have been deposited in GISAID under the accession numbers EPI_ISL_732948 - EPI_ISL_732959.

## Acknowledgements

We gratefully acknowledge the following Authors from the Originating laboratories responsible for obtaining the specimens and the Submitting laboratories where genetic sequence data were generated and shared via the GISAID Initiative, on which this research is partially based (Supplementary Table 1). MG thanks Alicja Rost, Ewa Zieliniewicz and Karolina Baranowicz for their assistance in the laboratory. LR thanks Bartosz Wasąg for helping in sequencing. We thank veterinary surgeons for their help in samples collection.

